# Incorporating in-source fragment information improves metabolite identification accuracy in untargeted LC-MS datasets

**DOI:** 10.1101/399105

**Authors:** Phillip M. Seitzer, Brian C. Searle

## Abstract

In-source fragmentation occurs as a byproduct of electrospray ionization. We find that ions produced as a result of in-source fragmentation often match fragment ions produced during MS/MS fragmentation and we take advantage of this phenomenon in a novel algorithm to analyze LC-MS metabolomics datasets. Our approach organizes co-eluting MS1 features into a single peak group and then identifies in-source fragments among co-eluting features using MS/MS spectral libraries. We tested our approach using previously published data of verified metabolites, and compared the results to features detected by other mainstream metabolomics tools. Our results indicate that considering in-source fragment information as a part of the identification process increases annotation quality, allowing us to leverage MS/MS data in spectrum libraries even if MS/MS scans were not collected.

## INTRODUCTION

Confidently identifying metabolites in LC-MS metabolomics datasets is a challenging problem^1^. Both targeted^2–3^ and untargeted^4^ LC-MS raw data can be internally or externally calibrated with chemical standards. While this can yield highly accurate metabolite detections, the approach is constrained to only measure endogenous levels of those standard metabolites. Additionally, external calibrant data must be reacquired when chromatographic conditions or instrument settings change, making it potentially prohibitively expensive and time-consuming to produce.

When internal or external standards are unavailable, metabolomics studies typically leverage several independent lines of evidence to detect metabolites, including accurate mass, retention time, and agreement between observed and theoretical isotopic peak intensities. Different types of identification information may be aggregated to produce a single identification score^5^, or identification probabilities using Bayesian networks^6^ and target-decoy approaches^7^. A popular alternative for analyzing untargeted LC-MS/MS data is matching acquired MS/MS against one or more large spectral libraries, such as NIST^8^, HMDB^9^, and METLIN^10^. While the number of features without MS/MS spectra acquired using data dependent acquisition (DDA) experiments remains significant, efforts to increase the number of MS1 features fragmented by the mass spectrometer^11^ and applications of data independent acquisition^12–13^ may improve data consistency.

However, many metabolomics experiments are still collected using LC-MS only, and even in LC-MS/MS datasets, many features only contain MS1 information. Without MS/MS information, search engines can only use accurate mass and isotopic distributions based on molecular formulae to detect metabolites^14^. As many metabolites share molecular formulae, scanning MS1-only data against spectral libraries yields incomplete, ambiguous, or partial metabolite identifications. Additionally, when individual metabolites ionize, they can produce unanticipated MS1 features as a result of neutral losses, in-source fragmentation, multimerization, and adducts^12,15^, further complicating the annotation process.

Here we present an approach to identify metabolites in untargeted LC-MS data by identifying in-source fragments that match to fragment peaks in MS/MS spectral libraries. To accomplish this, we have developed an algorithm to form consensus MS1 peak groups from a set of raw data files and use those peak groups in library searching. We have tested our method by comparing the feature detection, deisotoping and grouping steps of our algorithm to two mainstream open-source approaches using a complex LC-MS dataset containing 75 verified compounds. We find that our feature detection, deisotoping and peak grouping steps identify more of the verified compound features than other approaches. We also find that identifying in-source fragments in LC-MS data and including this information as a part of our identification process improves the accuracy of metabolite identifications.

## EXPERIMENTAL PROCEDURES

We downloaded mzML raw data from the Metabolights study 67 (MTBLS67)^16^ from the Metabolights raw data portal^17^. We processed raw files with MSConvertGUI (Proteowizard version 3.0.9987)^18^ to strip them of MS/MS scans, and generated both a centroided set and an uncentroided set of sample files (using the parameters cwt centroiding, snr = 0.1, peakSpace = 0.1).

We independently processed the uncentroided positive and negative mode files using Scaffold Elements 2.0.0 with search parameters that were chosen to match the original MTBLS67 study (specific search parameters are listed in **Supporting Information Table 1**). Monoisotopic peaks were searched against the NIST 2017^8^ and METLIN^10^ spectral libraries, as well as an empty library (to generate a baseline list of all detected features). We also generated an R script **(Supporting Information Script 1)** using Bioconductor^19^ to drive XCMS^20–21^ (version 3.0.2) and CAMERA^22^ (version 1.34.0). The script performed peak detection, peak grouping, and isotope detection on both the uncentroided sample files (using XCMS “matchedFilter”^20^) and the centroided sample files (using XCMS “centWave”^21^). We analyzed positive mode and negative mode files separately using search parameters that were chosen to match the original study (specific search parameters are listed in **Supporting Information Script 1**). The *m/z* and retention time coordinates of the 75 verified metabolites were compared to all monoisotopic *m/z* and retention time features identified by XCMS-matchedFilter + CAMERA, XCMS-centWave + CAMERA, and Scaffold Elements (script available in **Supporting Information Script 2**).

## RESULTS AND DISCUSSION

### The Scaffold Elements algorithmic workflow

We have developed an automated workflow to identify metabolites from untargeted LC-MS raw files using spectral libraries **(Fig 1a)**. Briefly (see **Supporting Information Note 1** and **Supporting Information Figure 1** for further details), we first organize raw data into isotopic feature clusters (IFCs) that contain a monoisotopic [M+0] feature, and [M+1] and [M+2] isotopic features. IFCs from the same sample are formed into MS1 peak groups based on elution profile **(Fig 1b)**. This step ensures that all ions produced during ionization of a single metabolite remain organized together. Failure to properly account for ionization effects can lead to ion misannotation, especially of in-source fragments^23^. We then align MS1 peak groups from all samples in the experiment to form cross-sample consensus MS1 peak groups. The formation of consensus elements is based on a number of independent metrics, including MS1 spectral similarity, peak shape, and agreement in *m/z* and retention time **(Fig 1c)**. Finally, we search consensus MS1 peak groups against spectral libraries and score metabolite groups and clusters **(Fig 1d)**. Score values increase both with agreement (higher mass accuracy and agreement with theoretically predicted isotopic distributions) and the amount of evidence associated with a metabolite annotation (number of ion types and in-source fragments identified).

**Figure 1.**
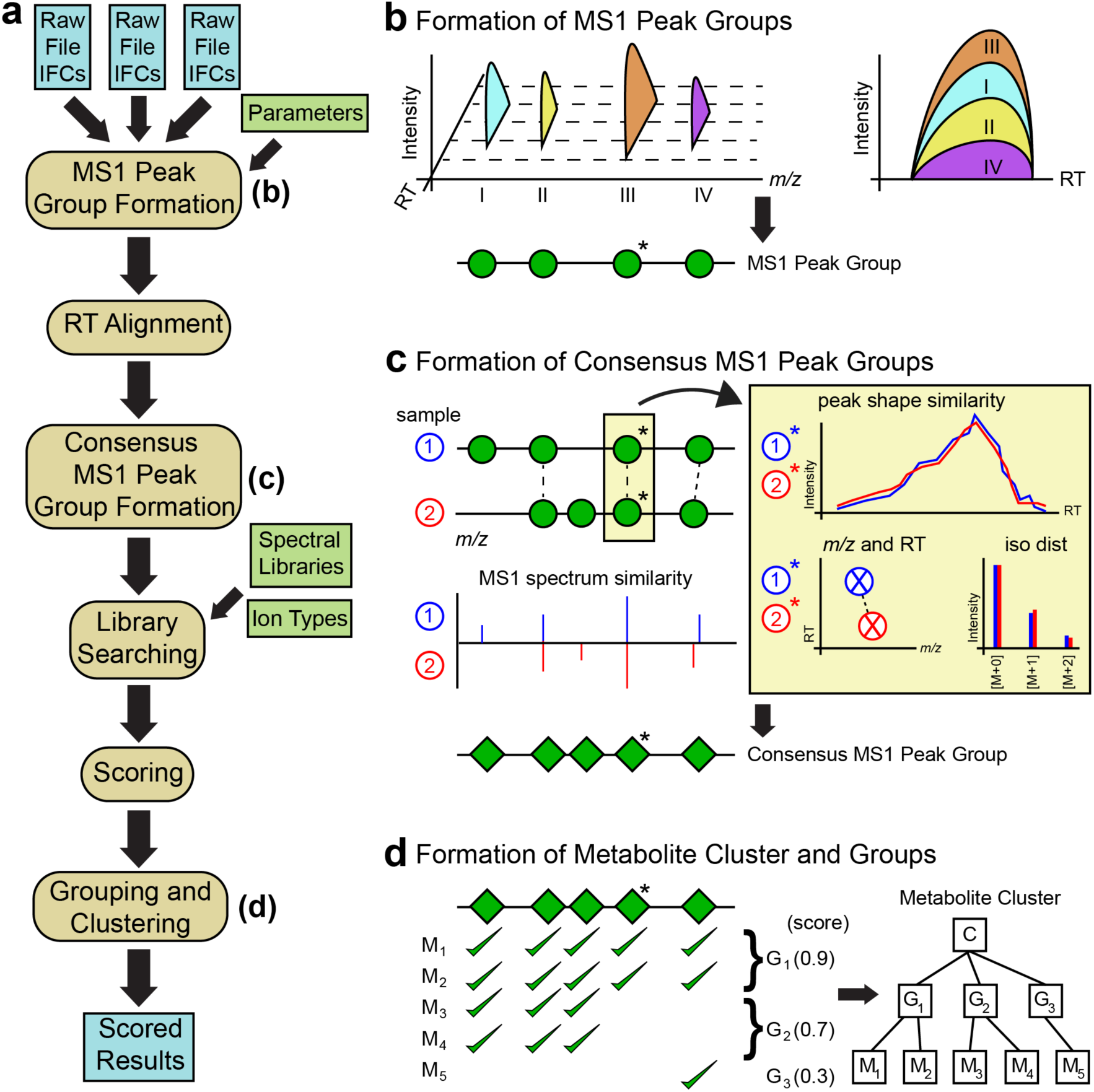
Scaffold Elements metabolite identification and scoring algorithm. **(a)** Complete workflow of Scaffold Elements identification and scoring algorithm. Tan, rounded boxes indicate algorithmic steps, green boxes indicate user-specified inputs, and blue boxes indicate algorithmic inputs and outputs. **(b)** An MS1 peak group is formed in a single sample by combining four co-eluting isotopic feature clusters (IFCs) (I, II, III, and IV). IFCs are represented as green circles on a line, with an asterisk indicating the most intense IFC in the peak group. **(c)** A consensus MS1 peak group is formed by comparing MS1 peak groups from each sample. A cross-sample MS1 spectrum similarity score is evaluated considering all IFCs in each peak group, and additional comparisons are made between a representative IFC from each MS1 peak group individually (light yellow boxes). The resulting consensus MS1 peak group is represented as green diamonds on a line, with an asterisk indicating the most intense consensus IFC in the consensus MS1 peak group. **(d)** Multiple putatively identified metabolites are organized into groups and clusters based on the consensus IFCs within a consensus MS1 peak group. In this schematic, a consensus MS1 spectrum of five IFCs was identified by five metabolites, which were organized into a cluster containing three groups, one of which contained only a single metabolite. Identification scores (shown next to each group in parentheses) indicate the most likely metabolite annotation for this cluster.

### Development of a “gold-standard” MS1-only dataset

We benchmarked our approach using the Metabolights study 67 (MTBLS67) ^16^. This study identified and quantified 75 yeast metabolites from nitrogen-starved *Saccharomyces Pombe* whole cell lysates using DDA-based LC-MS/MS. Sajiki et al confirmed the MS/MS fragmentation patterns and retention times of these metabolites using external standards. In an effort to produce a “gold-standard” MS1-only dataset of a complex metabolome with endogenous targets, we stripped these raw files of MS/MS scans. This produced a mock MS1-only data set containing 75 independently verified compounds.

### Comparing peak detection algorithms

We compared the peak detection, isotopic clustering, and peak grouping steps of our approach to two XCMS-based workflows, either XCMS “matchedFilter” ^20^ or XCMS “centWave” ^21^ peak detection, both followed by CAMERA isotopic grouping^22^. Scaffold Elements was executed without library matching to generate a list of all dataset features. We found that Scaffold Elements was able to detect more of the features associated with verified metabolites than either XCMS-CAMERA workflow, including 12 that were not identified by either approach **(Fig 2a)**. However, since Scaffold Elements reported more features than either XCMS-CAMERA workflow **(Supporting Information Figure 2)**, we were concerned that there would be a higher chance of noise matching a verified metabolite *m/z* and retention time coordinate by chance. To ensure that Scaffold Elements returned well-formed peaks, we manually investigated the features associated with the 12 metabolites that were only identified by Scaffold Elements. We found that 11 of these 12 verified metabolite features had a clear, reproducible signal **(Supporting Information Figure 3)**. Extracted ion chromatograms of features corresponding to one representative verified metabolite (Lysine) are shown in **Fig 2b**.

**Figure 2:**
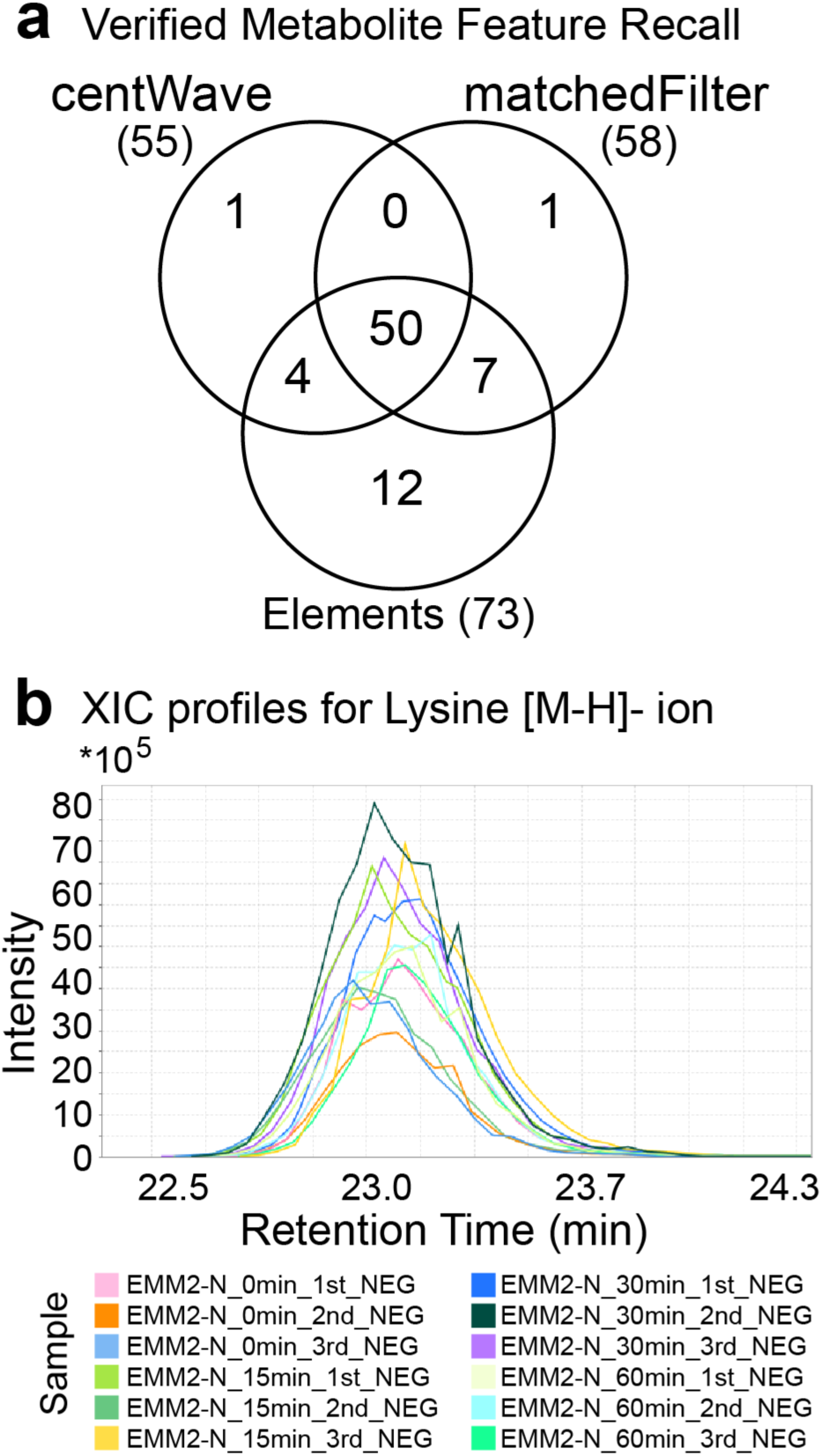
Scaffold Elements feature detection comparison. **(a)** Comparison of verified metabolite features identified by XCMS-centWave + CAMERA (centWave), XCMS-matchedFilter + CAMERA (matchedFilter) and Scaffold Elements (elements). Scaffold Elements identified 73 of the 75 features associated with verified metabolites, including 12 that were not detected by either XCMS-CAMERA workflow. **(b)** Extracted ion chromatograms (XICs) of a verified metabolite ion for Lysine ([M-H]-ion), which was identified only in Elements. The overlay plot of XICs shows a reasonable peak shape for this ion, which was independently identified in all 12 negative mode samples and correctly organized together into a single feature group.

### Using in-source fragments in scoring improves annotation quality

We next aimed to determine if searching for in-source fragments in MS1 peak groups improved metabolite annotation quality. We searched the MTBLS67 sample files with the NIST^8^ and METLIN^10^ spectral libraries, which together contained 65 of the 75 verified metabolites **(Supporting Information Table 2)**. Our feature detection algorithm identified the correct *m/z* and retention time feature for 63 of these 65 metabolites. However, multiple library annotations were returned for these features. Scaffold Elements’ scoring algorithm organized these annotations into clusters of metabolite groups, and ranked the annotations within each metabolite group.

We evaluated metabolite detection performance based on three metrics. For each independent search, we determined the proportion of correct annotations (where the annotation had the highest score in the metabolite group), unambiguous annotations (where the correct annotation had a uniquely higher score than all other annotations in the metabolite group), and unmistakable annotations (where the correct annotation was the only annotation in the metabolite group). Our approach of incorporating in-source fragment information in scoring improved all three of these metrics, notably increasing the proportion of unambiguous and unmistakable annotations by 22% and 60%, respectively **(Table 1)**. In many cases, the inclusion of in-source fragments in the search yielded rich MS1 peak groups that matched multiple MS/MS fragment peaks from the corresponding library spectrum with high mass accuracy **(Fig 3)**.

**Table 1.** Annotation of verified metabolites with and without consideration of in-source fragmentation (ISF) events in the identification process. ^a^The annotation had the highest score. ^b^The correct annotation had a uniquely higher score than all other annotations. ^c^The correct annotation was the only annotation in the metabolite group.

| Search | Correct<sup>a</sup> | Unambiguous<sup>b</sup> | Unmistakable<sup>c</sup> |
| --- | --- | --- | --- |
| **Major Ion** | 96.8% | 59.7% | 40.3% |
| **Major Ion + ISFs** | 98.4% | 72.6% | 64.5% |

**Figure 3:**
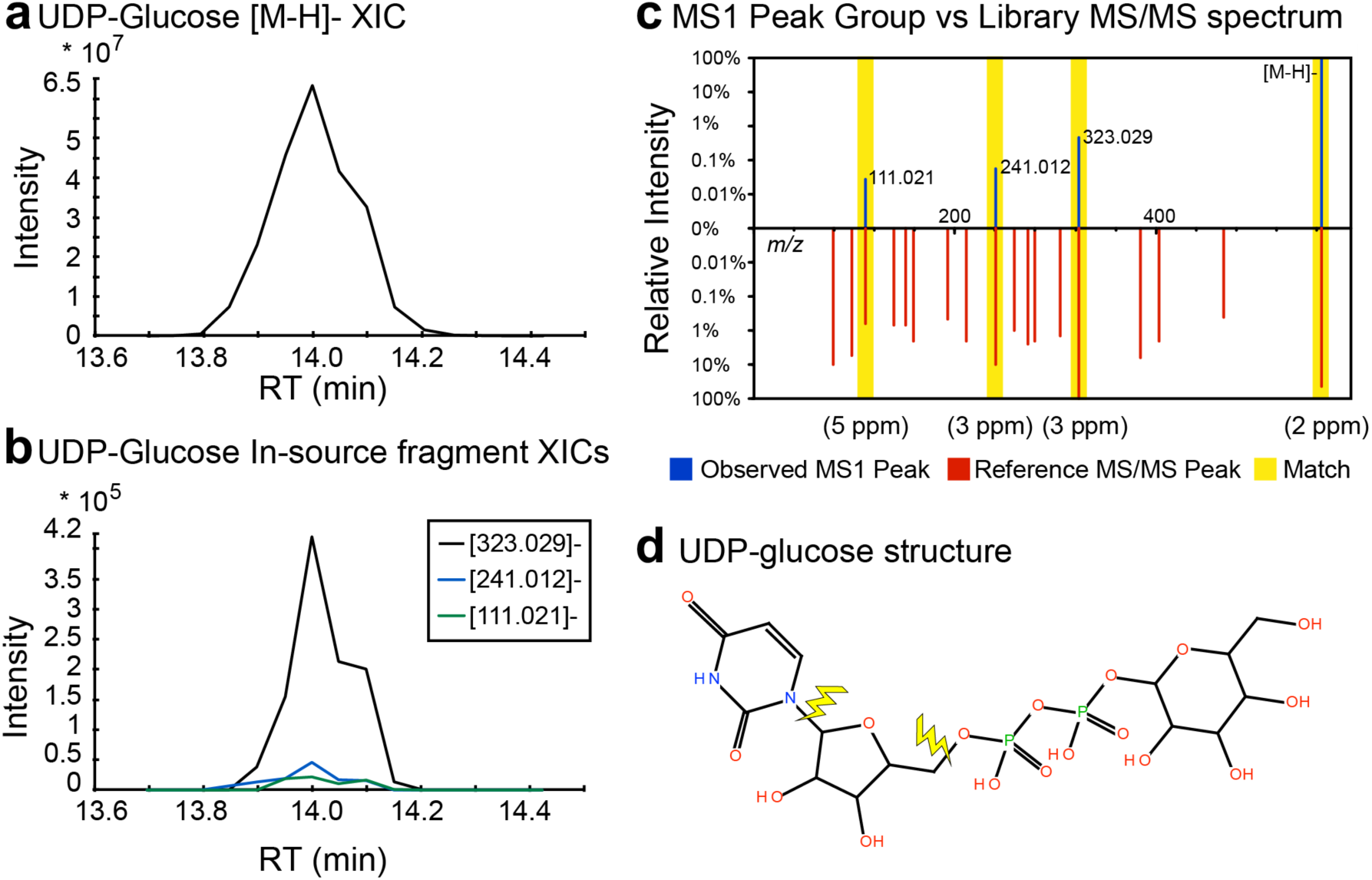
UDP-glucose MS1 peak group. An extracted ion chromatogram (XIC) of [M-H]-ion of UDP-glucose and **(b)** XICs of three detected in-source fragment peaks. **(c)** A butterfly plot comparing observed MS1 peak group of UDP-glucose ([M-H]-ion and three in-source fragment peaks) to METLIN library spectrum ID:6698 (METLIN ID 3598). Intensities are shown as a relative percentage to max spectral peak on a logarithmic scale to allow visualization of low-intensity peaks. The mass tolerance in ppm for each peak match is shown below butterfly plot. **(d)** The structure of UDP-glucose, with fragmentation sites corresponding to observed in-source fragments indicated by yellow lightning bolts. All observed data in figure was taken from the sample “EMM2-N_0min_2nd_NEG”.

## CONCLUSIONS

We have developed an approach to account for ionization effects by forming consensus MS1 peak groups prior to spectral library matching, and to use in-source fragments in those groups to perform pseudo-MS/MS library searching. Our results indicate that considering in-source fragments as part of the identification process improves confidence in metabolite detections. To increase the availability of these algorithms, we have made this tool available as a module in the Scaffold Elements software package distributed by Proteome Software.

Our results also demonstrate a caveat of spectral library search-based approaches: it is only possible to identify metabolites that are present in the specific spectral library (or libraries) searched. In our case, only 65 (86.7%) of the verified compounds were present in the NIST and METLIN spectral libraries **(Supporting Information Table 2)**. If a compound is present in the data but absent from the library, the compound will either be misidentified or remain unidentified. Without prior knowledge of which compounds are actually contained in the data, we can use our scoring approach to determine which annotations correspond to real compounds and which are misidentifications. We believe that improving candidate scoring is particularly important for analyzing untargeted metabolomics LC-MS data, as the ground truth identification might be absent from the library.

## [ASSOCIATED CONTENT]

**Supporting Information Note 1**: **Detailed description of Scaffold Elements 2.0 metabolite identification and scoring algorithm**. A detailed description of the Scaffold Elements 2.0 metabolite identification and scoring algorithm. Also includes a description of feature finding and isotopic grouping.

**Supporting Information Figure 1**: **Feature finding algorithm** Diagram of major steps of Scaffold Elements feature detection algorithm.

**Supporting Information Figure 2**: **Number of features identified by different programs** Summary of number of features identified by XCMS-centWave + CAMERA, XCMS-matchedFilter + CAMERA, and Scaffold Elements.

**Supporting Information Figure 3**: **XICs of verified features only detected by Scaffold Elements** Description and summary of 12 verified features only detected by Scaffold Elements, including overlaid XICs (showing XIC of each feature in all samples where it was detected).

**Supporting Information Table 1**: **Scaffold Elements parameters** Table of parameter used in all Scaffold Elements analyses.

**Supporting Information Table 2**: **Detailed Results of in-source fragment annotation comparison analysis** Detailed summary of the annotation results for 75 verified metabolites with and without consideration of in-source fragments.

**Supporting Information Script 1: XCMS CAMERA workflows (R script)**. R Script for generating (*m/z*, RT) feature list files using both XCMS-matchedFilter on profile mode files and XCMS-centWave on centroided files. Uses Bioconductor, XCMS, and CAMERA (for isotopic grouping).

**Supporting Information Script 2: Comparison of XCMS CAMERA workflows vs Scaffold Elements (Java script)** Java script comparing output of Scaffold Elements and XCMS-CAMERA workflows to features corresponding to verified metabolites.

## [AUTHOR INFORMATION]

### Corresponding Author

Brian C. Searle

### Author Contributions

The study was conceived by B.C.S. and P.M.S. The algorithm was implemented and evaluated by P.M.S. P.M.S. and B.C.S. wrote the paper. All authors have given approval to the final version of the manuscript.

## [ACKNOWLEDGEMENT]

We would like to acknowledge the entire staff at Proteome Software, Inc. for fruitful scientific discussions and feedback associated with development and implementation of the algorithm.

## [ABBREVIATIONS]

LC-MS: liquid chromatography mass spectrometry
LC-MS/MS: liquid chromatography tandem mass spectrometry
MS1: mass spectrometry
MS/MS: tandem mass spectrometry
IFC: isotopic feature cluster
NIST: national institute of standards and technology
HMDB: human metabolome database
MTBLS: Metabolights
DDA: data-dependent acquisition
DIA: data-independent acquisition
RT: retention time
ISF: in-source fragment
IFC: isotopic feature cluster
XIC: extracted ion chromatogram

